# Automatic Removal of False Connections in Diffusion MRI Tractography Using Topology-Informed Pruning (TIP)

**DOI:** 10.1101/338624

**Authors:** Fang-Cheng Yeh, Sandip Panesar, Jessica Barrios, David Fernandes, Kumar Abhinav, Antonio Meola, Juan C. Fernandez-Miranda

## Abstract

Diffusion MRI fiber tracking provides a non-invasive method for mapping the trajectories of human brain connections, but its false connection problem has been a major challenge. This study introduces topology-informed pruning (TIP), a method that automatically identifies singular tracts and eliminates them to improve the tracking accuracy. The accuracy of the tractography with and without TIP was evaluated by a team of 6 neuroanatomists in a blinded setting to examine whether TIP could improve the accuracy. The results showed that TIP improved the tracking accuracy by 11.93% in the single-shell scheme and by 3.47% in the grid scheme. The improvement is significantly different from a random pruning (p-value < 0.001). The diagnostic agreement between TIP and neuroanatomists was comparable to the agreement between neuroanatomists. The proposed TIP algorithm can be used to automatically clean up noisy fibers in deterministic tractography, with a potential to confirm the existence of a fiber connection in basic neuroanatomical studies or clinical neurosurgical planning.

## Introduction

Diffusion MRI fiber tracking provides a non-invasive way of mapping macroscopic connections in the human brain, but its accuracy issue still remains a challenge [1, 2]. A large-scale study examined 96 methods submitted from 20 research groups, showing that the accuracy of tractography quantified ranged from 3.75% to 92% due to differences in reconstruction methods and algorithms [3], and several strategies were proposed to improve fiber tracking, including motion correction, eddy current correction, and tractography clustering. Nonetheless, even if an optimal strategy demonstrated improvement, there were still a substantial amount of false connections [4].

To better understand the accuracy issue, our recent HCP-842 tractography study investigated false connections generated by deterministic tractography approaches [5]. By recruiting human neuroanatomists to analyze the neuroanatomical labels and to eliminate false continuities, the study identified two major causes for false connections: premature termination and false continuity. Premature terminations can be partly addressed by using a white matter mask. The mask allows for automatic checking of the fiber trajectory endpoints, and then false endpoints are rejected to achieve a better accuracy. Based on this paradigm, studies have previously used demarcation between white and gray matter boundaries derived from T1-weighted images as a tract-termination benchmark to cope with premature termination problem [6]. False tracts were determined as those prematurely terminating in white rather than gray matter, whilst real tracts were defined as those which terminated in gray matter, as determined by the overlap of T1 and diffusion-weighted images. However, delineating an accurate white matter mask can be complicated by image distortion in addition to the resolution mismatch between the diffusion-weighted and T1-weighted images. It is possible that this termination check may introduce another error due to an imperfect white matter mask.

The false continuity issue, on the other hand, is even more challenging. While the premature termination can be detected using a white matter mask, there is no effective strategy to detect false continuity using structural images or diffusion data. This is due to the limitation of diffusion MRI techniques in resolving the exact configuration of crossing fibers such as bending, fanning, or interdigitating [7]. Diffusion signals cannot alone differentiate these configurations within the voxel space. Until now, *a priori* knowledge of white matter anatomy was necessary to validate tractography results against the false continuity issue; nonetheless, our recent atlas study [5] demonstrated an interesting pattern that can potentially enable automated elimination of false tracks with false continuity: After separating whole-brain trajectories into clusters based on their neighboring distance, we found that clusters with fewer neighboring tracts were more likely to be false connections with false continuity. This finding suggests that a track with few neighboring tracts has a high likelihood of being a false continuity. One potential explanation of this phenomenon is that the false tracts arise between the overlapping boundaries of two real, adjacent fiber bundles. Since this overlapping boundary only forms a touching surface or line between two fiber pathways, it will only allow for a limited number of trajectories to pass through and cause false continuity. Moreover, it is known that errors of fiber tracking will accumulate during the tracking process [8]. As a result, the trajectories that pass through the overlapping boundaries tend to have very diverse propagation routes due to the perturbation around the boundaries. These two unique conditions combined will greatly reduce the chance of a false continuity tract to find a neighboring fiber trajectory. This observation thus brings forth our *singular tract hypothesis*—a singular tract (i.e. a trajectory with no neighboring tract) has a higher likelihood of being a false continuity connection. This hypothesis further inspired our topology-informed pruning (TIP) algorithm, which uses the topology of a tractogram itself to identify candidate false connections for removal. It was realized by first constructing a 3D tract density histogram to single out voxels with only one track passing through them, then subsequently eliminating those singular tracts to improve the accuracy. We examined the performance of TIP using data from our previous study [9], in which we tracked the arcuate fasciculus using a single region of interest (ROI) placed around the angular gyrus. The tractogram after TIP was then independently scrutinized by 6 neuroanatomists to assess whether TIP could reduce the percentage of false connections.

## Materials and Methods

### The TIP Algorithm

The TIP algorithm is illustrated in Fig.1. The input data are a set of trajectories obtained from the tractogram of a target fiber bundle (Fig.1a). The tractography of a fiber bundle can be obtained from a region-of-interest or any track selection approach. The number of trajectories should be high enough to produce a 3D histogram (yellow density map in Fig. 1b). A tract may pass through the same voxel more than one time, but it will only be counted once in the histogram. The second step is to identify voxels with a track count equal to one (red density map in Fig. 1b). These voxels are nearby white matter regions where singular tracts “go astray” via an overlapping boundary. These tracts are identified and excluded from the bundle (Fig. 1c) to produce a pruned tractogram (Fig. 1d). The whole process can be repeated until no more singular tracts are found. The computation complexity of TIP is O(*N*), where *N* is the number of trajectories. This TIP algorithm is fully automatic and requires no manual intervention to eliminate false connections.

**Fig. 1.**
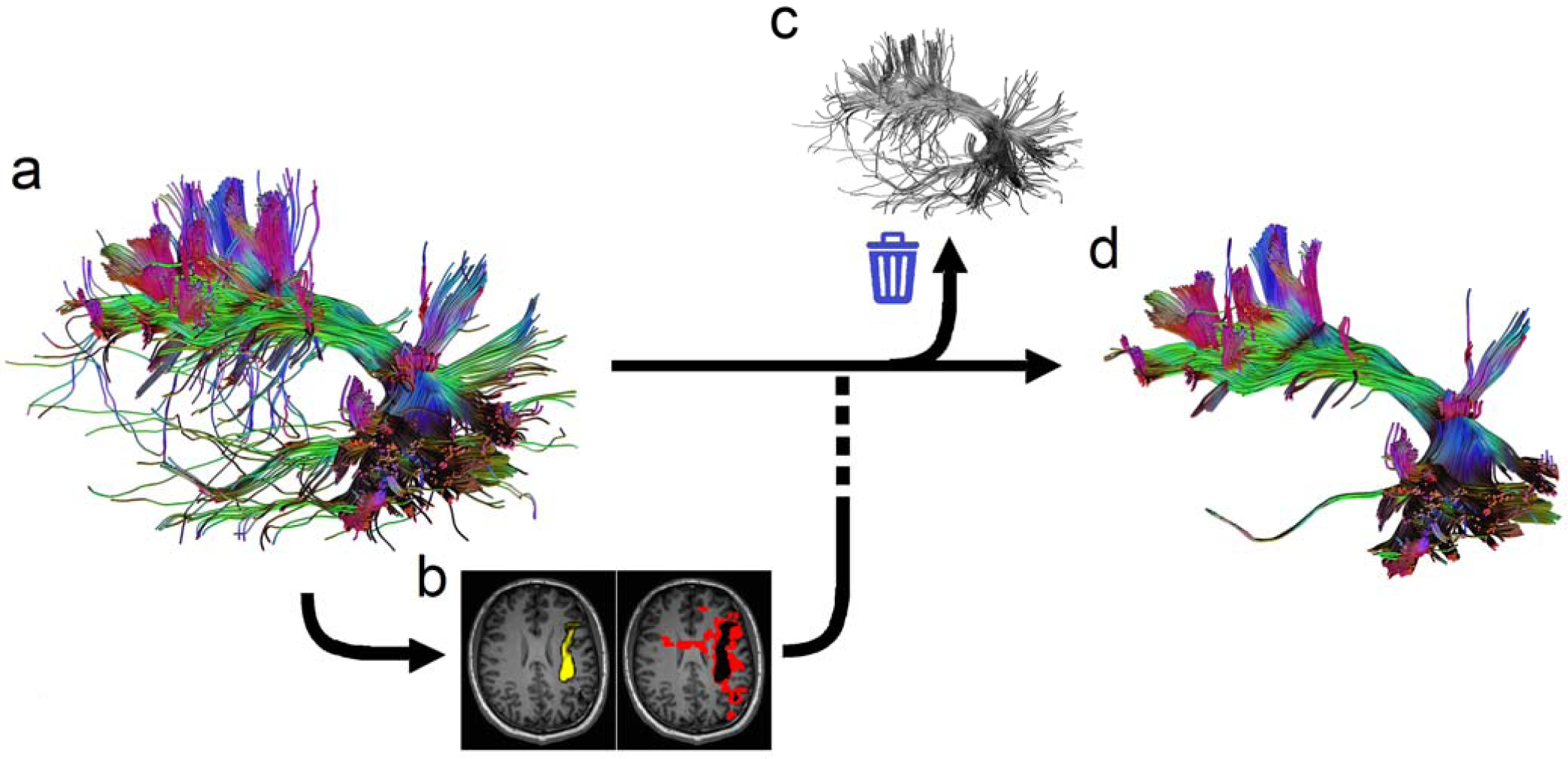
The topology-informed pruning (TIP) algorithm. (a) the input tractogram of a target fiber bundle is generated using the adequate density of seedings. (b) the tractogram is used to compute the density map (yellow color) for identifying voxels with low track density (red color). (c) Tracts passing low-density voxels are eliminated from the tractogram. (d) The resulting tractogram from the TIP algorithm eliminates singular tracts that are likely to be false connections.

### TIP applied to the arcuate fasciculus tractogram

We applied TIP to existing data from our previous tractography study [9] to examine whether TIP can improve the accuracy of tractography. The data were originally acquired from a normal subject scanned in a 3T Siemens Tim Trio MRI scanner, using a twice-refocused echo planar imaging (EPI) diffusion pulse sequence. Both a 253-direction single-shell scheme and a 203-direction grid scheme were acquired. For the 253-direction single shell scheme, the diffusion parameters were TR/TE = 7200 ms/133 ms, b-value = 4,000 s/mm^2^, and the scanning time was approximately 30 minutes. For the 203-direction grid scheme, the diffusion parameters were TR/TE = 7200 ms/144 ms, the maximum b-value = 4,000 s/mm2, and the scanning time was approximately 25 minutes. The shell and grid sampling schemes shared the same spatial parameters: the field of view was 240 mm × 240 mm, the matrix size was 96 × 96, and the slice thickness was 2.5 mm. A total of 40 slices were collected.

DSI Studio (http://dsi-studio.labsolver.org) was used to track the arcuate fasciculus using a method identical to the original study [9]. To summarize, the generalized q-sampling imaging reconstruction [10] was applied to both sampling schemes with a diffusion sampling length ratio of 1.25. A 14-mm radius spherical ROI was placed at the angular gyrus, and tracts passing the ROI connecting temporal with frontal regions were selected. The tractogram was generated using an angular threshold of 60 degrees and a step size of 1.25 mm. Whole-brain seeding was conducted until a total of 2,000 fiber tracts were generated from the ROI.

6 neuroanatomists (S.P., D.F., J.B., J.F., K.A., A.M.) independently examined the tractogram to identify false tracts. The examination was conducted by a blinded setting. They were instructed to remove false tracts from the tractogram and were unaware of the TIP algorithm, nor did they have access to the TIP-processed tractogram, or teach others results. The accuracy of the tractogram was then quantified using the following formula:

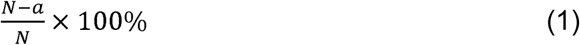

where *N* is the total number of tracts, and *a* is the number of tracts identified as false connections. The accuracy of the TIP-processed tractogram was calculated using the same formula, where *N* represents non-singular tracts, and *a* is the number of non-singular false tracts as identified by the neuroanatomists.

TIP could also be viewed as an observer, and we further quantified the diagnostic agreement between TIP and any of the neuroanatomists by the following formula:

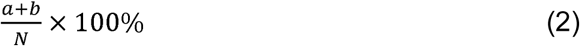

where *a* is the number of positive agreements quantified by the number of singular, false tracts simultaneously identified by TIP and the neuroanatomists. *b* is the number of negative agreements quantified by TIP as non-singular and “not-false” by the neuroanatomists. *N* is the number of total tracts evaluated. 6 diagnostic agreements were calculated between TIP and any of the 6 neuroanatomists. The diagnostic agreements between the neuroanatomists were also calculated for a comparison.

### TIP applied to peritumoral tractogram

We applied TIP to a 60-year-old female patient diagnosed with glioblastoma. The patient was scanned on a Siemens 3T Tim Trio scanner using a twice-refocused spin-echo diffusion sequence to acquire the diffusion data. A total of 514 diffusion directions were sampled with a maximum b-value of 7000 s/mm2. The in-plane resolution was 2.4 mm. The slice thickness was 2.4 mm. The diffusion data were reconstructed using generalized q-sampling imaging [10] with a diffusion sampling length ratio of 1.25. The restricted diffusion was quantified using restricted diffusion imaging [11] to differentiate between tumor regions, edematous area, and normal white matter tissue. The tumor region and peritumoral edematous area were manually delineated to facilitate tracking the peritumoral fiber pathways. Peritumoral tractogram was generated using a deterministic fiber tracking algorithm [9] with a region of interest placed at the peritumoral edematous region. A total of 10000 tracts were calculated, and the tractograms with and without TIP were compared to examine whether TIP could improve quality.

## Results

Figure 2a shows the tractogram of arcuate fasciculus before and after being pruned by the TIP algorithm. The tractogram generated using the angular gyrus ROI shows “noisy” fibers (Fig. 2a), especially using the single-shell scheme. The tractogram from the grid sampling scheme appears to be cleaner, but deviant trajectories are still visible. TIP effectively removes noisy fiber while retaining the main topology structure. The improvement is particularly striking in the single-shell scheme. The pruned tractogram appears to be consistent with the microdissection results (Fig. 2b, adapted from [9]).

**Fig. 2.**
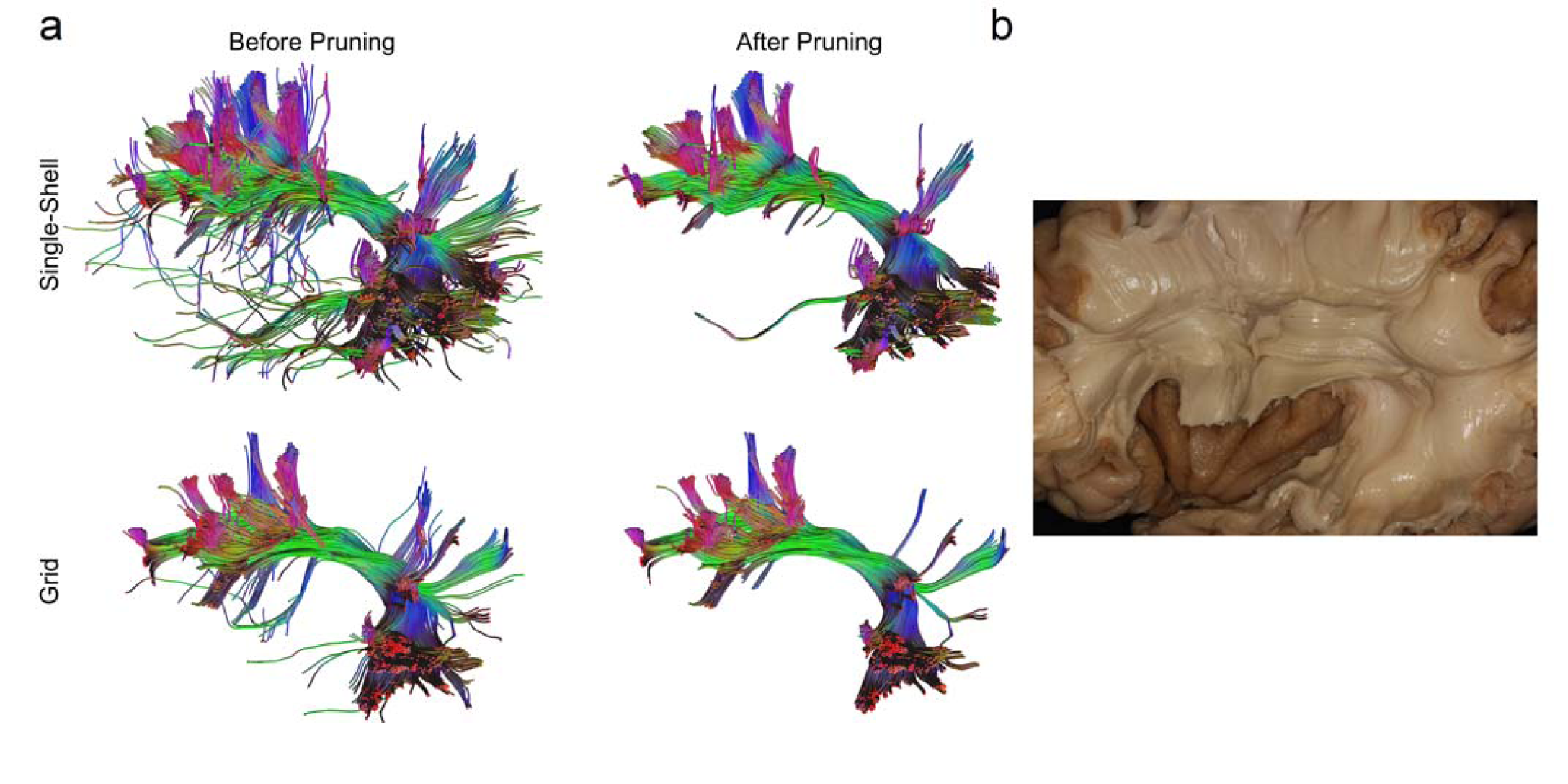
(a) The arcuate fasciculus tractogram after topology-informed pruning shows a coherent architecture with reduced noisy fibers. (d) Cadaver dissection of the nearby structures shows consistent structure (adapted from [9]).

### The accuracy of TIP-processed tractogram

The accuracy of tractography before and after TIP is quantified in Fig.3. The tractogram from a single-shell scheme is shown in Fig. 3a, whereas the tractogram from the grid scheme is shown in Fig. 3b. A total of 6 different evaluations were conducted independently by the 6 neuroanatomists. The lines connect evaluations from the same neuroanatomists. In both Fig. 3a and 3b, all evaluations unanimously showed improved accuracy after TIP was applied to the arcuate fasciculus tractography. The average improvement in the single-shell scheme was 11.93%, whereas the average improvement in the grid scheme was 3.47%. Improvement is most obvious in the single-shell dataset. The lower improvement in the grid scheme could be due to its lower sensitivity for demonstrating branching fibers, as shown in the original tractogram before pruning.

**Fig. 3.**
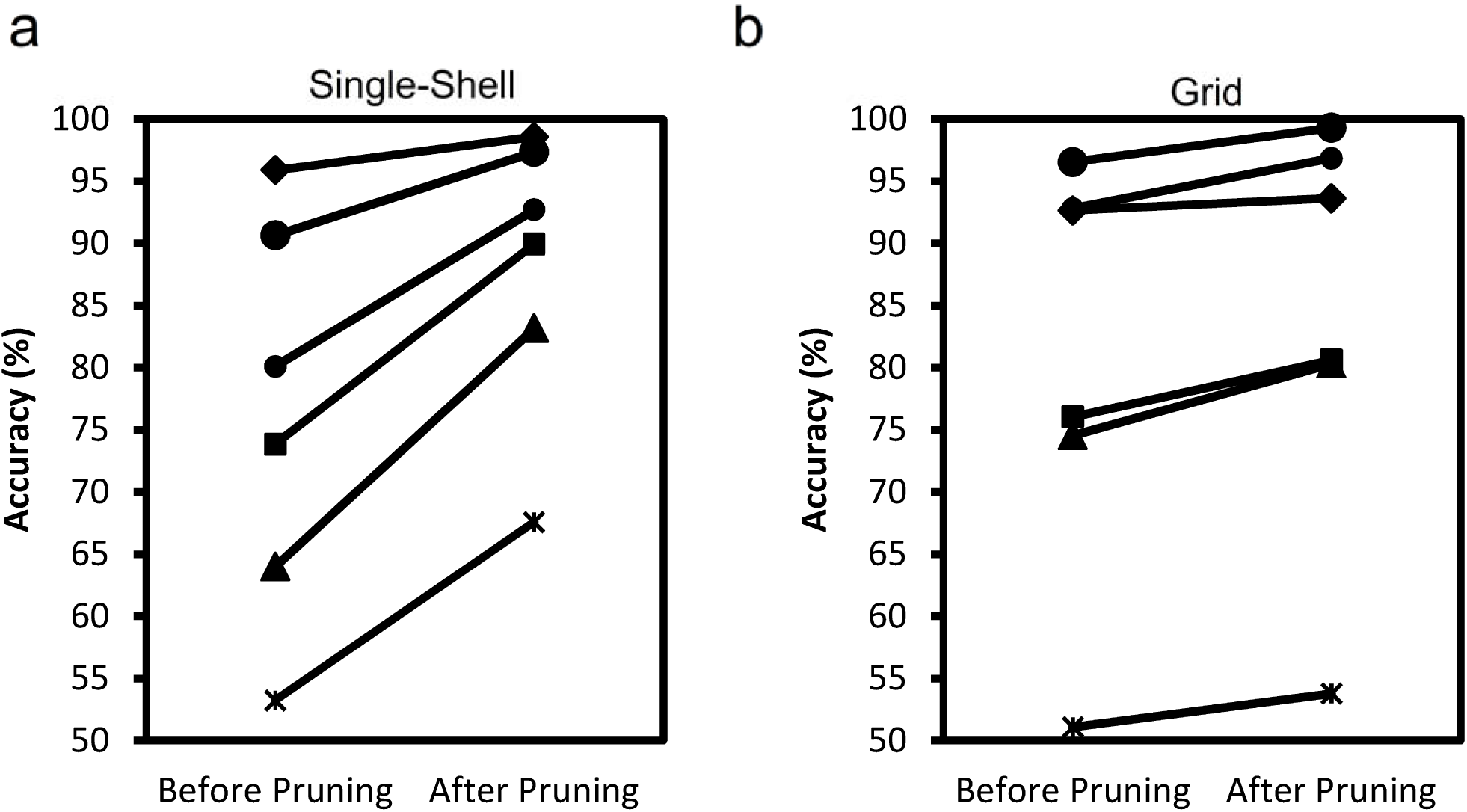
Line plots showing the accuracy of tractography before and after pruning in (a) single-shell data and (b) grid data. The line connects the two evaluations conducted by the same neuroanatomists. All evaluations show increased accuracy after pruning. The improvement is most prominent in the single shell dataset.

We further compare the performance of TIP with random pruning. Since random pruning (e.g. random elimination of an arbitrary number of tracts) will result in an equal opportunity between increased accuracy and decreased accuracy, the chance that 6 evaluations unanimously present improved accuracy in random pruning is thus 1/2^6^=0.015625 (follows a binomial distribution), and the probability of this happening on both single-shell and grid datasets is thus (1/2^6^)^2^ = 0.00024414062. Therefore, for a null hypothesis that TIP is no different from random pruning, the p-value will be smaller than 0.001 based on this frequentist hypothesis-testing framework, and thus we can conclude that TIP is significantly different from random pruning.

### Diagnostic agreement between TIP and neuroanatomists

Figure 4 compares the tractograms produced when false tracts are removed by a representative neuroanatomist, compared with the same tracts pruned by TIP. The neuroanatomist chosen here had the highest diagnostic agreement with the other 5 neuroanatomists and thus can be viewed as the representative neuroanatomist of the group. In both the single-shell and grid schemes, the TIP-processed tractogram was highly consistent with the neuroanatomist-pruned tractogram, though differences can still be observed at minor branches.

**Fig. 4.**
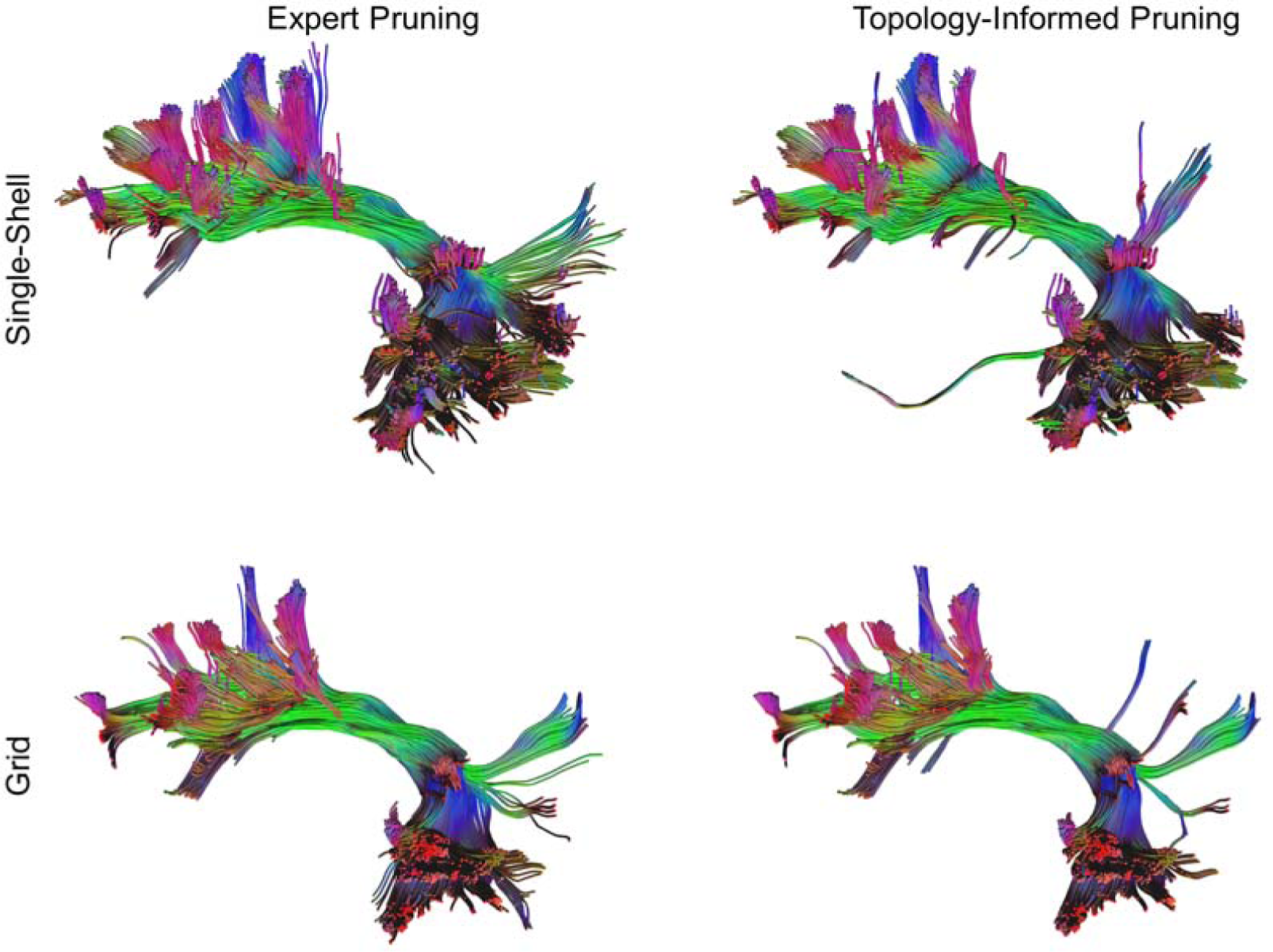
Tractogram pruned by a representative neuroanatomist (left) compared with the same tractogram pruned by TIP (right). Although minor differences can still be found, the overall results show remarkable consistency in the main structures and major branches.

We further evaluated the agreement of TIP with other neuroanatomists. Further quantification using diagnostic agreement showed that the averaged diagnostic agreement between TIP and 6 neuroanatomists was 77.33% with the single-shell scheme and 77.73% with the grid scheme. In comparison, the averaged diagnostic agreement between neuroanatomists was 77.46%±1.62% (standard error) with the single-shell scheme and 75.80%±2.14% with the grid scheme. The agreement between TIP and neuroanatomists was within the standard error of the mean agreement between the neuroanatomists. With the grid scheme, the average agreement between TIP and neuroanatomists was even slightly higher than the average between the 6 neuroanatomists. We can conclude that the agreement between TIP and neuroanatomists was comparable to the agreement within neuroanatomists.

### Removal of false connections in peritumoral tractogram

Figure 5 shows how TIP can be used to remove false connections in peritumoral tractography. The T1-weighted image is shown at the neurology convention in Fig. 5. The 3D reconstruction shows the tumor (flesh color) located at frontoparietal region and its peritumoral edematous area (light-blue color). Figure 5b shows the peritumoral tractogram generated directly from the fiber tracking algorithm after placing the region of interest at the peritumoral edematous region. The red arrows point to possible false connections that cross the several sulci due to false continuity. These pathways are false connections because there should be no pathway crossing a sulcus between two nearby gyri. Figure 5c shows the same tractogram processed automatically by two consecutive TIP runs to eliminate false connections. The false connections pointed by red arrows were eliminated without any manual invention, suggesting that TIP is an effective tool for improving the accuracy of diffusion MRI tractography.

**Fig. 5.**
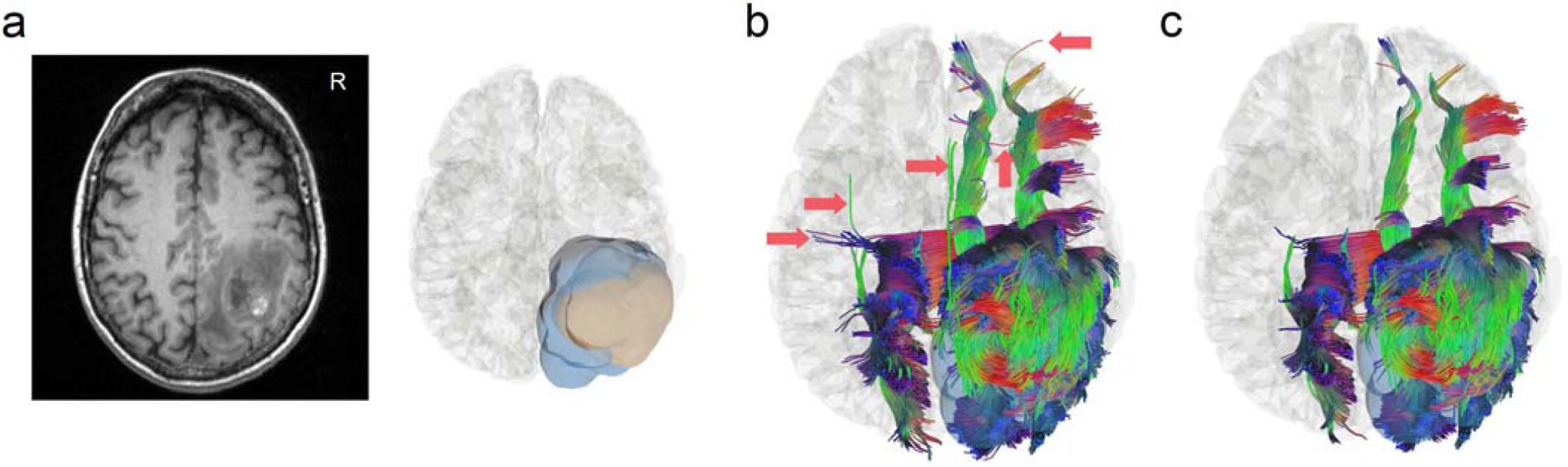
False connections in peritumoral tractogram removed by TIP. (a) The T1-weighted image shows a glioblastoma located at the right frontoparietal lobe of a 50-year-old patient, whereas the 3D reconstruction illustrates the extent of tumor (flesh-colored) and its surrounding edematous tissue (blue-colored). (b) Peritumoral tractogram generated from fiber tracking contains possible false connections annotated by red arrows. (c) TIP automatically removes most of these false connections without manual intervention.

## Discussion

Here we show that the topology of a tractogram can be used to prune it to achieve a better accuracy. The TIP algorithm reduced the percentage of false tracts by 11.93% in the single-shell scheme and 3.47% in the grid scheme. The performance difference can be explained by the fact that the single-shell tractography in our study captured more branching fibers, and the trajectories appeared to be “nosier”.The pruning effect was thus more dramatic in such a high sensitivity setting. Although the improvement with the grid scheme is limited, the evaluations by all 6 neuroanatomists unanimously indicated improved accuracy after pruning. Moreover, the agreement of TIP with neuroanatomists was comparable to the agreement between the neuroanatomists alone. This suggests that TIP can replace a substantial part of the evaluation work done by neuroanatomists to scrutinize tractography. We also demonstrated the real-world application of TIP on a tumor patient. The function was implemented as a shortcut in DSI Studio (Ctrl+T), which allowed the users to quickly remove false connections when inspecting peritumoral tractogram.

TIP appears to be a unique tract validation method using only the relationship between individual tracts within the tractogram as a benchmark. Its theoretical basis differs from other post-processing methods which rely on diffusion MRI signals, T1W, or other imaging modalities to remove false connections [6, 12]. While the T1W or other imaging modalities can be used to improve the accuracy, using them to capture false connections may introduce additional problems. For example, signal intensity in the T1W images can be inhomogeneous, and segmenting white matter regions from T1W adds further complexity. Using diffusion MRI signals as the prior may also present pitfalls of signal-distortion and artifacts in DWI. These methods view each track as an independent target to be evaluated and do not use topological information of the entire tractogram to eliminate false connections. The unique theoretical basis of TIP allows it to potentially complement other anatomical-prior methods to improve tracking accuracy.

The computation time for TIP in our study was negligible in DSI Studio relative to the time taken for 3D tractographic visualization. TIP only has linear complexity proportional to the track count (i.e. O(N)) and can be readily applied to tractograms with any number of trajectories. In comparison, a typical tract clustering algorithm would require O(N^2^), which has limited capability for application to large fiber-count bundles.

There are still limitations with TIP. We cannot rule out the possibility that a singular tract identified by TIP is a real connection. This adverse result occurs if a small branch of fiber bundles receives only limited seeding points due to its small size. Insufficient seeding can result in limited numbers of neighboring fibers, resulting in a valid connection being eliminated by TIP. The removal of these tracts results in reduced sensitivity of tractography to capture small fiber pathways or branches, meaning that the accuracy improvement from TIP comes with a trade-off for sensitivity to small branching fibers. Therefore, it is noteworthy that applying TIP to small pathways may have a risk of over pruning.

Moreover, applying TIP to whole brain tractography may not be an ideal utilization. The low-density voxels of a fiber bundle can be occupied by nearby pathways, making the detection of singular tracts difficult for TIP. A more sophisticated topology algorithm computing the distance between pair-wise tracts for identification of singular tracts may be more appropriate.

Changing parameter settings for the TIP algorithm could expand its potential applications. For example, the threshold for defining low-density voxels can be adjusted to adapt to a different seeding density. A high threshold will yield highly confirmative results to justify the existence of a fiber pathway. As more and more fiber tracking studies have been proposed to discover human brain pathways and their segmentation [13-16], TIP can strengthen the results by boosting the accuracy of tractography. The number of recursive iterations can be limited to a small number to allow for different pruning effects. For studies correlating structural connectivity with neuropsychological measures, a single iteration of TIP can complement group connectometry analysis to achieve better false discovery rates. In neurosurgical applications, TIP can assist mapping of peritumoral pathways to help neurosurgeons organize a detailed surgical plan.

In conclusion, the topology information from a fiber bundle can be used to identify possible false connections in order to achieve better anatomical validity. The TIP algorithm can be integrated with deterministic fiber tracking to map human brain connections with a higher accuracy.

